# Biphasic roles of hedgehog signaling in the production and self-renewal of outer radial glia in the ferret cerebral cortex

**DOI:** 10.1101/2021.04.12.439562

**Authors:** Shirui Hou, Wan-Ling Ho, Lei Wang, Bryan Kuo, Jun Young Park, Young-Goo Han

## Abstract

The neocortex, the center for higher brain function, emerged in mammals and expanded in the course of evolution. The expansion of outer radial glia (oRGs) and intermediate progenitor cells (IPCs) plays key roles in the expansion and consequential folding of the neocortex. Therefore, understanding the mechanisms of oRG and IPC expansion is important for understanding neocortical development and evolution. By using mice and human cerebral organoids, we previously revealed that hedgehog (HH) signaling expands oRGs and IPCs. Nevertheless, it remained to be determined whether HH signaling expanded oRGs and IPCs *in vivo* in gyrencephalic species, in which oRGs and IPCs are naturally expanded. Here, we show that HH signaling is necessary and sufficient to expand oRGs and IPCs in ferrets, a gyrencephalic species, through conserved cellular mechanisms. HH signaling increases oRG-producing division modes of ventricular radial glia (vRGs), oRG self-renewal, and IPC proliferation. Notably, HH signaling affects vRG division modes only in an early restricted phase before superficial-layer neuron production peaks. Beyond this restricted phase, HH signaling promotes oRG self-renewal. Thus, HH signaling expands oRGs and IPCs in two distinct but continuous phases during cortical development.

## Introduction

The neocortex, a defining feature of the mammalian brain, underwent robust evolutionary expansion culminating in the highly convoluted brains (gyrencephaly) of certain mammals. The emergence and expansion of the neocortex underlie sophisticated perception, cognition, and behavior. Neocortical expansion reflects an increased number of neural cells and, therefore, depends on increased populations and proliferation capacity of neural progenitor cells (NPCs) (Rakic 2009; Lui et al. 2011; Borrell and Götz 2014; Florio and Huttner 2014; Sun and Hevner 2014; Dehay et al. 2015). The primary NPCs are ventricular radial glia (vRGs), whose cell bodies form the ventricular zone (VZ) that surrounds the brain ventricle on the apical side of the developing brain; hence, they are also called apical radial glia (Malatesta et al. 2000; Noctor et al. 2001). vRGs are bipolar, with a short apical process contacting the brain ventricle and a long basal process contacting the pial surface, which latter radially guides the newly generated excitatory neuron through the intermediate zone to the cortical plate. vRGs can produce neurons directly or indirectly via secondary NPCs, including intermediate progenitor cells (IPCs) and outer radial glia (oRGs) (Haubensak et al. 2004; Miyata et al. 2004; Noctor et al. 2004; Fietz et al. 2010; Hansen et al. 2010; Reillo et al. 2011). IPCs and oRGs that are generated from vRGs translocate basally and form the subventricular zone (SVZ). Accordingly, oRGs are also called basal radial glia, intermediate radial glia, or translocating radial glia.

The SVZ and the NPCs therein are greatly expanded in gyrencephalic mammals, as compared with species that have small and smooth (lissencephalic) brains; consequently, they are thought to play important roles in neocortical expansion and folding. The expanded SVZ in gyrencephalic mammals can be divided into the inner SVZ (ISVZ) and the outer SVZ (OSVZ). The ISVZ is formed by a dense population of IPCs that are generated from vRGs. The OSVZ, which was first identified in primates (Smart et al. 2002; Lukaszewicz et al. 2005; Zecevic et al. 2005), forms later than and superficial to the ISVZ. The OSVZ contains oRGs, which are detached from the ventricular surface, and IPCs generated from oRGs. Whereas the ISVZ is similar to the SVZ in lissencephalic mammals, the OSVZ and oRGs are relatively specific to gyrencephalic mammals, with some notable exceptions. For example, the common marmoset, a near-lissencephalic primate, has an enlarged OSVZ with abundant oRGs (Garcia-Moreno et al. 2012; Kelava et al. 2012), indicating that an enlarged OSVZ and oRGs are not sufficient to induce gyrencephaly. Gyrification occurs after neurons are produced; therefore, developmental processes beyond increased neuron production, such as neuronal differentiation and gliogenesis, might also be important for the development of convoluted brains (Hevner and Haydar 2012; Kroenke and Bayly 2018; Rash et al. 2019). Although the OSVZ and oRGs are not sufficient for the development of gyrencephaly, the expansion of the OSVZ and oRGs is thought to enable the expansion and maintenance of NPCs beyond the spatially limited neurogenic area surrounding the brain ventricle. Like vRGs, oRGs have a prolonged proliferative potential and a basal process, which increase the production and dispersion of neurons, contributing to an expanded and convoluted neocortex (Fietz et al. 2010; Hansen et al. 2010; Reillo et al. 2011). Therefore, elucidating the mechanisms that regulate oRGs is fundamental to understanding neocortical development and evolution.

Hedgehog (HH) signaling is an important regulator of nervous system development from neural progenitor specification and proliferation to circuit formation (Garcia et al. 2018). Secreted HH proteins (Sonic, Indian, and Desert HH in mammals) bind to Patched1 (PTCH1), a 12-transmembrane receptor that inhibits the 7-transmembrane protein Smoothened (SMO). Binding of HH to PTCH1 relieves this inhibition, enabling activation of SMO and its localization to the primary cilium. Activated SMO in the primary cilium inhibits Suppressor of Fused (SUFU)–facilitated formation of repressor forms of GLI-family zinc-finger transcription factors (GLI2 and GLI3) and promotes GLI activator formation, leading to the expression of target genes. Notably, mutations that decrease HH signaling activity, such as duplication of *PTCH1* or inactivating mutations in *SMO*, can cause microcephaly (Nanni et al. 1999; Heussler et al. 2002; Ginocchio et al. 2008; Derwińska et al. 2009), whereas mutations that increase HH signaling activity, such as activating mutations in *SMO* or inactivating mutations in *PTCH1* or *SUFU*, can cause megalencephaly in humans (Twigg et al. 2016; Shiohama et al. 2017; Klein et al. 2019). In mice, HH signaling is also necessary and sufficient to expand the neocortex (Komada et al. 2008; Wang et al. 2016). Together, these findings suggest that HH signaling is a conserved mechanism that promotes neocortical growth.

Recently, we showed that HH signaling is necessary and sufficient to expand both oRGs and IPCs in mice (Wang et al. 2016). HH signaling expands oRGs by changing vRG division modes to produce oRGs and by increasing oRG self-renewal. HH signaling expands IPCs by increasing their self-amplifying divisions. Whether these cellular functions of HH signaling in mice are conserved in gyrencephalic species is an important question that must be answered in order to understand the development and evolution of gyrencephalic brains. We have addressed this question by using the ferret, a gyrencephalic carnivore, as a model system. The ferret has been an important model for studying the developmental mechanisms of gyrencephalic brains because it has a distinct OSVZ with abundant oRGs and, compared to mice, has a relatively longer neurogenic period that continues after birth (Jackson et al. 1989; Noctor et al. 1997; Fietz et al. 2010; Reillo et al. 2011; Martínez-Cerdeño et al. 2012; Reillo and Borrell 2012; Kawasaki 2014). By using pharmacologic manipulation of HH signaling, we found that HH signaling changes vRG division modes, oRG self-renewal, and IPC proliferation in ferrets as it does in mice. Notably, we also found that HH signaling can change vRG division modes during only a limited period of neocortical development in both mice and ferrets. The results of our studies using ferrets show that HH signaling is a conserved and key factor promoting oRG and IPC expansion and neocortical growth in both lissencephalic and gyrencephalic mammals.

## Materials and Methods

### Animals

Timed pregnant ferrets (*Mustela putorius furo)* were purchased from Marshall BioResources and kept on a 12-h light / 12-h dark cycle in the Animal Resource Center at St. Jude Children’s Research Hospital (St. Jude). All procedures were performed according to a protocol approved by the Institutional Animal Care and Use Committee at St. Jude. We treated pregnant ferrets with vismodegib (25 mg/kg/day; 5 mg/mL stock in 0.5% hydroxypropylmethylcellulose/0.2% Tween 20) or SAG (25 mg/kg/day; 5 mg/mL stock in 0.5% methylcellulose/0.2% Tween 20) by oral gavage after subcutaneously injecting a mild tranquilizer (acepromazine, 0.2–0.5 mg/kg). We collected embryos by cesarean section from pregnant ferrets anesthetized with isoflurane. After surgery, ferrets were euthanized with a barbiturate overdose. To collect postnatal brains, ferret kits were anesthetized with avertine (400–500 mg/kg) and isoflurane and perfused with 4% paraformaldehyde.

### Cortical Slice Culture

We maintained cortical slices in culture by following a published procedure (Gertz et al. 2014). Embryonic brains were dissected out in ice-cold artificial cerebrospinal fluid (aCSF) containing 125 mM NaCl, 2.5 mM KCl, 1 mM MgCl_2_, 2 mM CaCl_2_, 1.25 mM NaH_2_PO_4_, 25 mM NaHCO_3_, and 25 mM D-(+)-glucose and embedded in 3.5% low-melting-point agarose in aSCF. Brain slices of 350–500 µm thickness were sectioned into ice-cold aCSF by using a vibrating microtome (Leica VT1200 S), transferred to 0.45-µm Millicell-CM culture inserts (MilliporeSigma, cat. no. PICM03050), and pre-incubated in slice culture medium containing 66% Basal Medium Eagle (BME) (Gibco, cat. no. 21010046), 25% Hanks’ Balanced Salt Solution (HBSS) (Gibco, cat. no. 14170112), 5% (v/v) fetal bovine serum (FBS), 2 mM GlutaMAX (Gibco, cat. no. 35050061), 1% (v/v) N2 Supplement (Gibco, cat. no. 17502048), and 1% (v/v) penicillin/streptomycin.

Vismodegib (200 nM), SAG (200 nM), or DMSO was added to the culture medium daily. The slices were then fixed in 4% formaldehyde (pre-chilled to 4°C) for 1 h and processed as cryopreserved samples.

### Immunohistochemistry

The brains were immersed in 30% sucrose in phosphate-buffered saline (PBS) overnight then embedded in OCT medium (Sakura Finetek). Tissue blocks were cryosectioned at a thickness of 16 μm, and the sections were transferred to glass slides (Superfrost Plus Microscope Slides, Fisher Scientific, cat. no. 12-550-15). For immunolabeling, the cryosections were warmed to room temperature (RT) and subjected to heat-induced epitope retrieval (HIER) in 0.1% citrate buffer, pH 6.0, with 1% glycerol in a steamer (Oster 5712 Electronic 2-Tier 6.1-Quart Food Steamer) for 16 min. After cooling to RT, the slides were washed three times in PBS for 5 min each and permeabilized in 0.3% Triton X-100 in PBS for at least 1 h. Primary antibodies were diluted in blocking solution containing 5% Normal Donkey Serum (Sigma-Millipore, cat. no. D9663) and 0.1% Triton X-100 in PBS and incubated overnight at 4°C. After an immediate rinse with PBS plus 0.1% Triton X-100, the slides were washed three more times for 10 min each. Secondary antibodies were diluted 1:200 in blocking solution and incubated for at least 2 h at RT. Coverslips were mounted with Aqua-Poly/Mount (Polysciences, Inc.), and the slides were stored at 4°C. Primary antibodies and their dilutions were as follows: sheep-anti-BrdU (abcam ab1893), 1:500; chicken anti-GFP (Novus Biologicals, cat. no. NB100-1614), 1:500 to 1:1000; mouse anti–phospho-histone H3 (phospho Ser10) (abcam, cat. no. 14955), 1:500; rabbit anti–phospho-histone H3 (Ser10) (Millipore-Sigma, cat. no. 06-570), 1:2000; mouse anti-KI67(MIB1, DAKO, M7240), 1:100; rabbit anti-PAX6 (clone: Poly19013; BioLegend, cat. no. 901301 [previously Covance, cat. no. PRB-278P]), 1:200 to 1:300; sheep anti-PAX6 (AF8150: R&D Systems, 5 µg/mL); rat anti-TBR2 (clone: Dan11mag; eBioscience, Fisher Scientific, cat. no. 50-245-556), 1:200 to 1:250; mouse anti-SATB2 (Santa Cruz Biotechnology, sc-81376), 1:50; rat anti-CTIP2 (abcam, ab18465), 1:500; chicken anti-TBR1 (Sigma-Millipore, AB2261), 1:200; and mouse anti– phosphorylated vimentin (Ser55) mAb (MBL Life Science, cat. no. D076-3), 1:1000. Alexa Fluor 488-, 568-, and 647-conjugated donkey anti-rabbit, anti-rat, anti-mouse, anti-sheep, and anti-chicken secondary antibodies were purchased from Invitrogen, Abcam, and Jackson ImmunoResearch Inc.

### Microscopy, Image Processing, and Quantification

Fluorescence confocal images were acquired with a Zeiss 780 confocal microscope. Maximal projections of four or five Z-sections, 0.5 to 1 µm apart, were made using the image-processing module in the Zen Black software (Zeiss). The exported maximal projection images were analyzed using Adobe Photoshop and ImageJ software. We quantified marker-expressing progenitor cells in radial columns perpendicular to the ventricular surface. The column density of each progenitor cell type was obtained by dividing the number of cells in the column by the width of the column base at the ventricular surface. The column densities were normalized to the vehicle-treated controls for each experiment. The average value for the control samples was set as 100%.

### Analysis of Modes of vRG Division

Brain sections were labeled with antibodies to phosphorylated vimentin (P-VIM) and histone H3 phosphorylated at Ser10 (PH3) and stained with 4’,6-diamidino-2-phenylindole (DAPI). Cells at anaphase or telophase were used to determine the modes of division. The angle formed by their cleavage plane and apical surface, termed alpha (α), was measured using ImageJ. The modes of vRG division were defined as horizontal divisions (60 ≤ α ≤ 90), oblique divisions (30 ≤ α < 60), or vertical divisions (0 ≤ α < 30).

### Statistical Analysis

Statistical analysis was performed using GraphPad Prism software. The chi-square test was used to compare the distribution of the angles of cell divisions. The Mann–Whitney test was used to compare the populations of NPCs. Three animals per group were used.

## Results

### HH Signaling is Necessary and Sufficient to Expand IPCs and oRGs in the Ferret Cortex

To examine whether HH signaling regulated NPCs in the neocortex in gyrencephalic species *in vivo*, we orally treated pregnant ferrets with vismodegib, a specific inhibitor of HH signaling, at embryonic days 33 (E33) and 34 (E34) (25 mg/kg/day) and collected embryonic brains at E35 (Fig. 1A). The neurogenesis stage in the cortex of E33–E35 ferrets corresponds to the E14 stage in the mouse cortex (Workman et al. 2013). We chose to use E33–E35 because HH signaling is necessary and sufficient to expand IPCs and oRGs from E13.5 in mice without disrupting the patterning of NPCs (Yabut et al. 2015; Wang et al. 2016). We chose vismodegib because it penetrates brains (Robinson et al. 2015) and is approved by the US Food and Drug Administration for treating HH signaling–driven tumors through oral delivery. We determined the vismodegib dose by extrapolating from the doses used for rat embryos (Morinello et al. 2014). Vismodegib treatment at E33 and E34 did not grossly affect the size and morphology of embryos (not shown) or the histology and vasculature of their brains (Supplementary Fig. 1). HH signaling is a well-established mitogen for granule neuron precursor cells (GNPs) in the cerebellum. Therefore, to ensure the effectiveness of the vismodegib treatment, we examined the proliferation of GNPs in the cerebellum. As expected, inhibition of HH signaling via vismodegib strongly decreased the proliferation of GNPs *in vivo* (Supplementary Fig. 1).

**Fig. 1.**
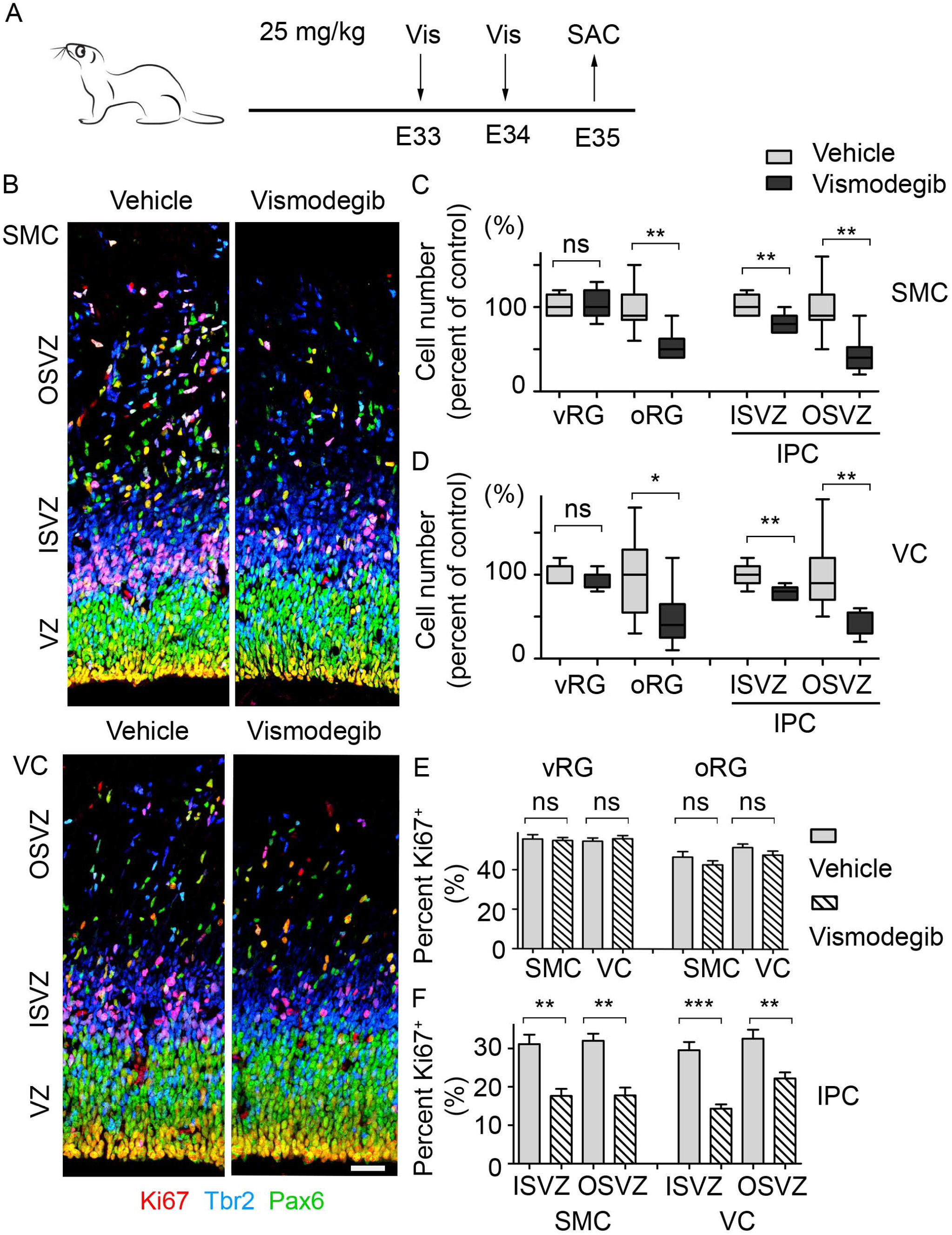
HH signaling is required for oRG and IPC expansion in the sensorimotor cortex (SMC) and the visual cortex (VC). (*A*) Diagram of the treatment strategy. Pregnant ferrets were treated with vismodegib (Vis), a HH signaling inhibitor, or with vehicle (Ve) by oral gavage at E33 and E34. Embryonic brains were collected at E35. (*B*) Micrographs of the E35 SMC and VC labeled with antibodies to Pax6 (green), Tbr2 (blue), and Ki67 (red). Scale bar = 50 μm. (*C, D*) Box plots with min/max whiskers that show the relative numbers of vRGs, oRGs, and IPCs. (*E, F*) Quantification of the proliferating NPCs that expressed Ki67. Mean ± SEM. Mann–Whitney test: ns, *P* > 0.05; * *P* < 0.05; ** *P* < 0.005; *** *P* < 0.001.

Next, we examined whether vismodegib affected NPCs in the cerebral cortex. Vismodegib significantly decreased the number of oRGs (Pax6^+^ Tbr2^−^ cells located above the VZ) and IPCs (Tbr2^+^ cells) without affecting the number of vRGs (Pax6^+^ Tbr2^−^ cells located in the VZ) in both the rostral (sensorimotor cortex) and caudal (visual cortex) areas of the cortex (Fig. 1). To understand the mechanism by which vismodegib decreased the numbers of oRGs and IPCs, we quantified apoptotic cells and proliferating cells by staining for an apoptotic marker, cleaved caspase 3, and a proliferation marker, Ki67. Vismodegib did not change the number of apoptotic cells (Supplementary Fig. 2). It significantly decreased the proportion of proliferating IPCs but did not affect the proportions of proliferating vRGs or oRGs (Fig. 1E and F). The decrease in Ki67^+^ Tbr2^+^ IPCs also suggested that the inhibition of HH signaling increased cell cycle exit of Tbr2^+^ IPCs. Together, these results show that HH signaling is necessary to expand oRGs and IPCs and to promote the proliferation of IPCs during E33–E35 in ferrets.

To test whether HH signaling was sufficient to expand NPCs in the cerebral cortex, we orally treated pregnant ferrets with SAG, a specific activator of HH signaling, at E33 and E34 (25 mg/kg/day) and collected embryonic brains at E35. SAG treatment significantly increased the number of oRGs and IPCs, but not that of vRGs (Fig. 2A–C). Furthermore, SAG selectively increased the proliferation of IPCs (Fig. 2D and E). Thus, the augmented activation of HH signaling induced effects that were the exact opposite of those induced by the inhibition of HH signaling. Taken together, these results show that HH signaling is necessary and sufficient to expand oRGs and IPCs and to promote the proliferation of IPCs in ferrets during E33–E35 *in vivo*.

**Fig. 2.**
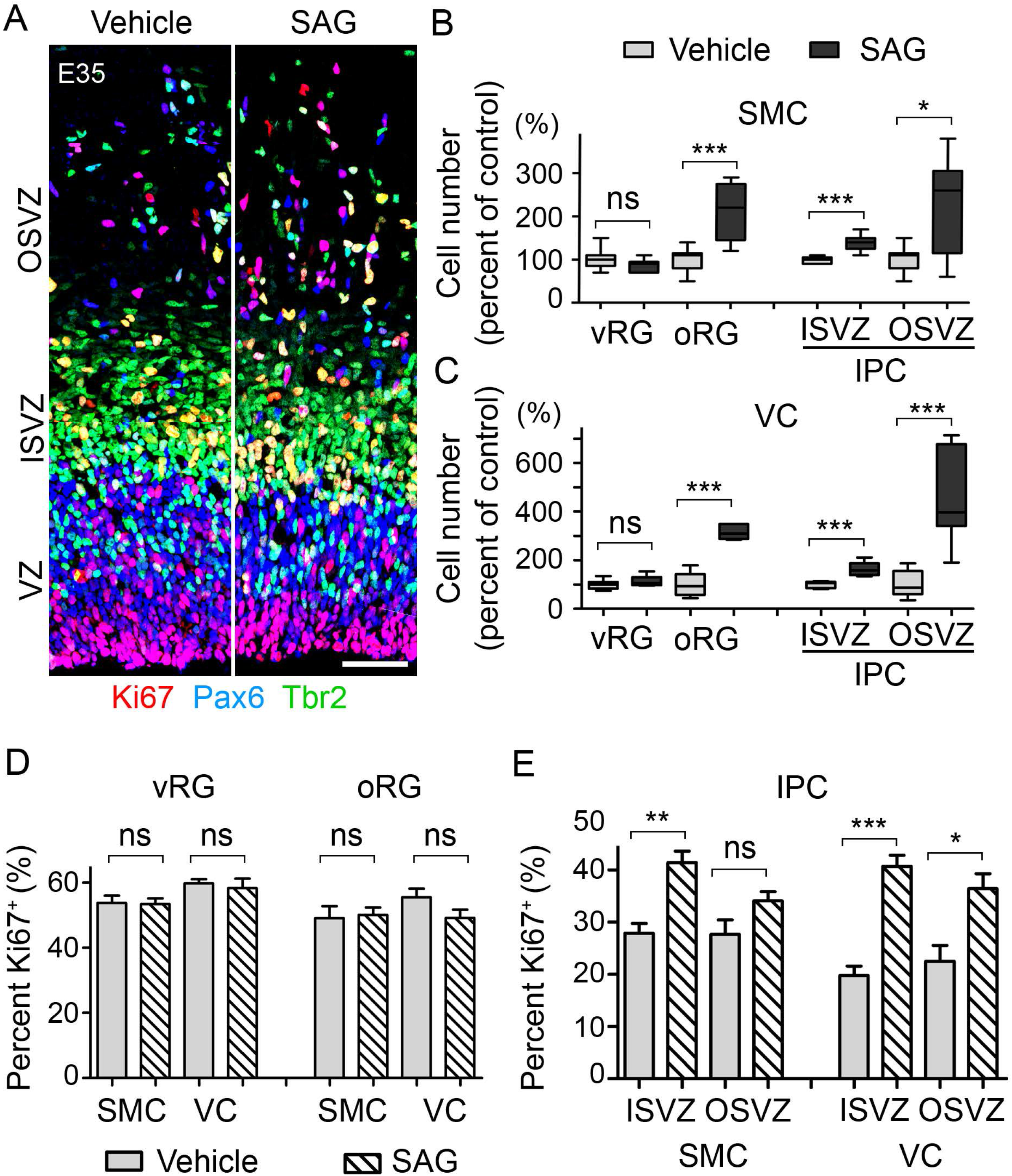
HH signaling is sufficient for oRG and IPC expansion. (*A*) Micrograph of E35 visual cortex labeled with antibodies to Pax6 (green), Tbr2 (blue), and Ki67 (red) after treatment with SAG, an HH signaling activator, at 25 mg/kg or vehicle (Ve) for 2 days (E33 and E34). Scale bar = 50 μm. (*B, C*) Box plots with min/max whiskers that show the relative numbers of vRGs, oRGs, and IPCs in the sensorimotor cortex (SMC) and the visual cortex (VC). (*D, E*) Quantification of the proliferating NPCs that expressed Ki67 in the SMC and the VC. Mean ± SEM. Mann–Whitney test: ns, *P* > 0.05; * *P* < 0.05; ** *P* < 0.01; *** *P* < 0.001.

### HH Signaling Changes the Division Angle of vRGs in a Limited Developmental Window

Our results indicated that HH signaling increased the number of oRGs through mechanisms that were independent of increasing their proliferation. Previous live-imaging studies had shown that the division angle of a vRG relative to the ventricular surface is highly associated with the fate of its daughter cells (Shitamukai et al. 2011; LaMonica et al. 2013). vRGs dividing on an axis horizontal to the ventricular surface mostly produce neurons or IPCs, whereas those dividing obliquely or vertically (collectively called non-horizontally dividing vRGs) produce oRGs. Therefore, to test whether HH signaling promoted oRG production in ferrets, we examined the division angles of the vRGs. To determine the division angles, we identified mitotic vRGs in E35 cortices by immunostaining for phosphorylated histone H3 (PH3) and phosphorylated vimentin (P-Vim) and staining with DAPI (Fig. 3A). At E35, 62% of vRGs in the visual cortex divided obliquely or vertically in vehicle-treated embryos (Fig. 3B). Remarkably, inhibition of HH signaling by vismodegib significantly decreased non-horizontal divisions to 33% (Fig. 3B), suggesting that HH signaling promotes non-horizontal, oRG-producing vRG divisions in the visual cortex at E35. In sharp contrast to vRGs in the visual cortex, only 17% of vRGs in the sensorimotor cortex of vehicle-treated embryos divided non-horizontally, and vismodegib treatment failed to reduce this fraction further (Fig. 3C). The high proportion of non-horizontal vRG divisions in the visual cortex at E35 is consistent with the findings of a previous study that, in ferrets, vRGs generate oRGs destined for the OSVZ during a restricted period of cortical development, with a burst between E34 and E36 in the visual cortex (Martínez-Martínez et al. 2016). Like other mammalian cerebral cortices, the ferret cerebral cortex develops along a rostral-to-caudal gradient, with the visual cortex developing later than the somatosensory cortex (Jackson et al. 1989; Noctor *et al*. 1997). In ferrets, germinal zones also develop in the same rostral-to-caudal direction; at E34, the OSVZ is already apparent in the rostral cortex but not in the visual cortex (Reillo and Borrell 2012). Consistently, the OSVZ was thicker at E35, with many more cells in the sensorimotor cortex than in the visual cortex (Fig. 1B). Therefore, the small proportion of non-horizontal vRG divisions and the finding that vismodegib had no effect on the vRG division angle in the sensorimotor cortex suggest that most vRGs in the sensorimotor cortex have already exited the developmental window to produce oRGs by E35 and that endogenous HH signaling outside this window does not affect vRG division angles. These findings also suggest that vismodegib reduced the number of oRGs in the sensorimotor cortex through a mechanism other than altering the vRG division angle.

**Fig. 3.**
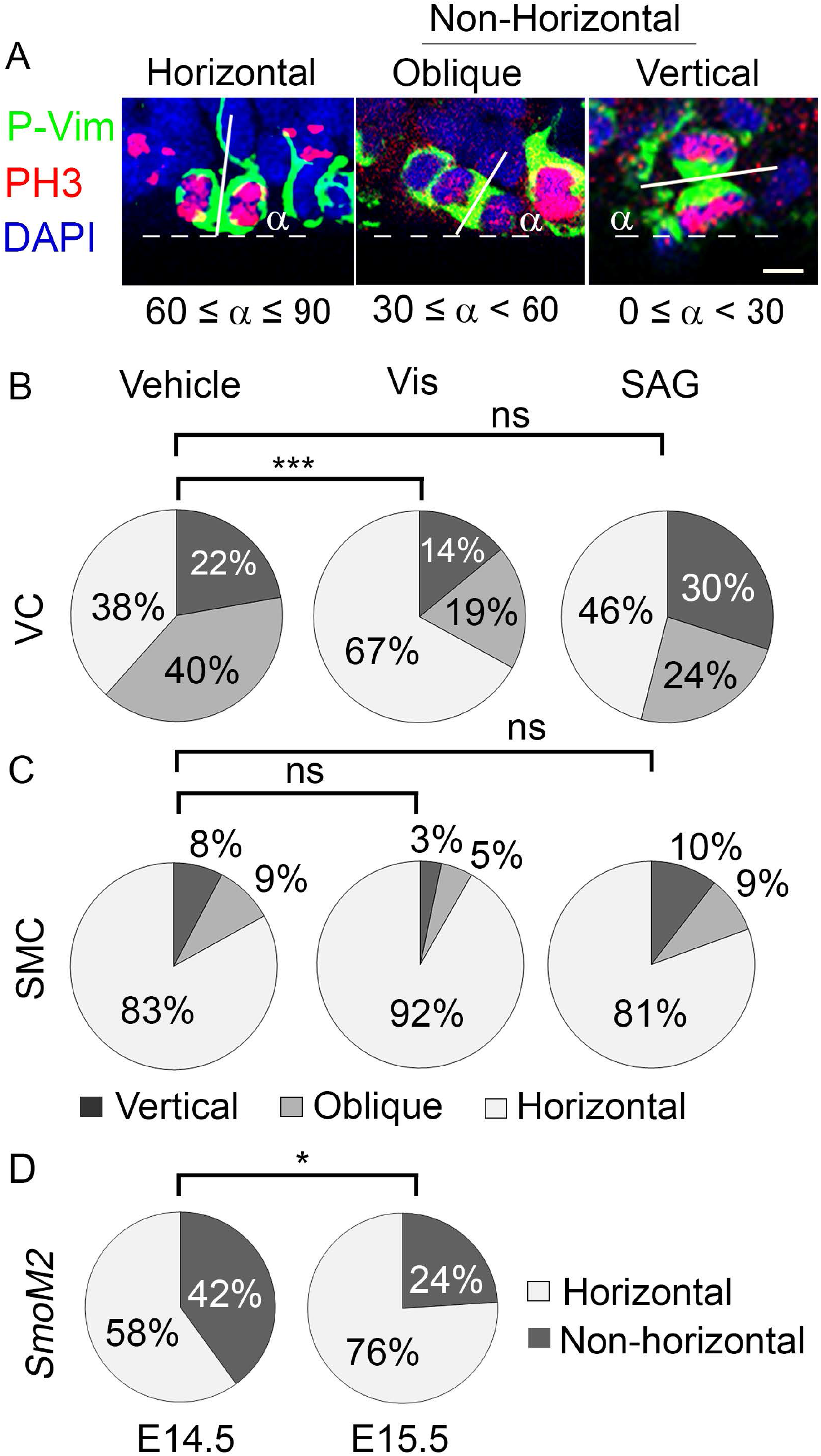
HH signaling promotes non-horizontal division of vRGs. (*A*) Micrographs of E35 vRGs labeled with antibodies to phospho-vimentin (P-Vim) (green) and phospho-histone H3 (PH3, Ser10) (red) and stained with DAPI (blue). Scale bar = 5 μm. Solid lines indicate the cleavage planes of anaphase cells, and dotted lines indicate the ventricular surface. α indicates the acute angle between the solid and dotted lines. The modes of division are categorized as “horizontal” (60° < α ≤ 90°), “oblique” (30° ≤ α < 60°), or “vertical” (0° ≤ α < 30°), with oblique and vertical divisions being collectively referred to as “non-horizontal” divisions. (*B*, C) Quantification of the acute angle (α) in the visual cortex (VC) (*B*) and the sensorimotor cortex (SMC) (*C*). (*D*) Quantification of horizontal vs. non-horizontal divisions of vRGs in *GFAP::Cre*; *SmoM2* mice at E14.5 and E15.5. The E14.5 values are those reported by Wang et al. (2016) and are shown to enable direct comparison. Chi-square test: ns, *P* > 0.05; * *P* < 0.05; *** *P* < 0.001.

Next, we asked whether increased HH signaling could affect vRG division angles. In contrast to vismodegib treatment, increasing HH signaling with SAG treatment did not change the vRG division angles (Fig. 3B and C). Note that the same SAG treatment did increase the number of oRGs and IPCs (Fig. 2). These results suggest that the effect of intrinsic HH signaling on the vRG division angle is already at a maximum at this stage. To test this possibility, we prepared cultures of embryonic brain slices from E33 embryos and treated them with SAG (200 nM), vismodegib (200 nM), or DMSO for 2 days *ex vivo* (Fig. 4). We reasoned that the endogenous source of HH in a brain slice of 350–500 µm thickness might not produce enough HH to affect the vRGs in a brain slice exposed in excess culture medium, thereby enabling us to test directly whether the activation of HH signaling by an added activator affected vRG division angles. Consistent with our reasoning, the basal level of non-horizontal vRG divisions in the visual cortex was decreased *ex vivo* relative to that *in vivo* (41% *ex vivo* vs. 62% *in vivo, P* < 0.005) (Figs. 3B and 4C) and vismodegib failed to decrease non-horizontal vRG divisions further *ex vivo*. (Fig. 4C). In contrast to the lack of effect of vismodegib *ex vivo* and SAG *in vivo*, SAG significantly increased non-horizontal divisions in both visual and sensorimotor cortices *ex vivo* (Fig. 4C). Therefore, *ex vivo*, there was no innate difference between the vRGs in those two cortical areas with respect to their response to HH signaling, which was sufficient to promote non-horizontal vRG divisions. We prepared brain slices at E33, when vRGs in both the sensorimotor and the visual cortices may be in the window of oRG production. E33 vRGs in the sensorimotor cortex might have maintained their competency to produce oRGs in culture, allowing an HH signaling activator to change the vRG division angles.

**Fig. 4.**
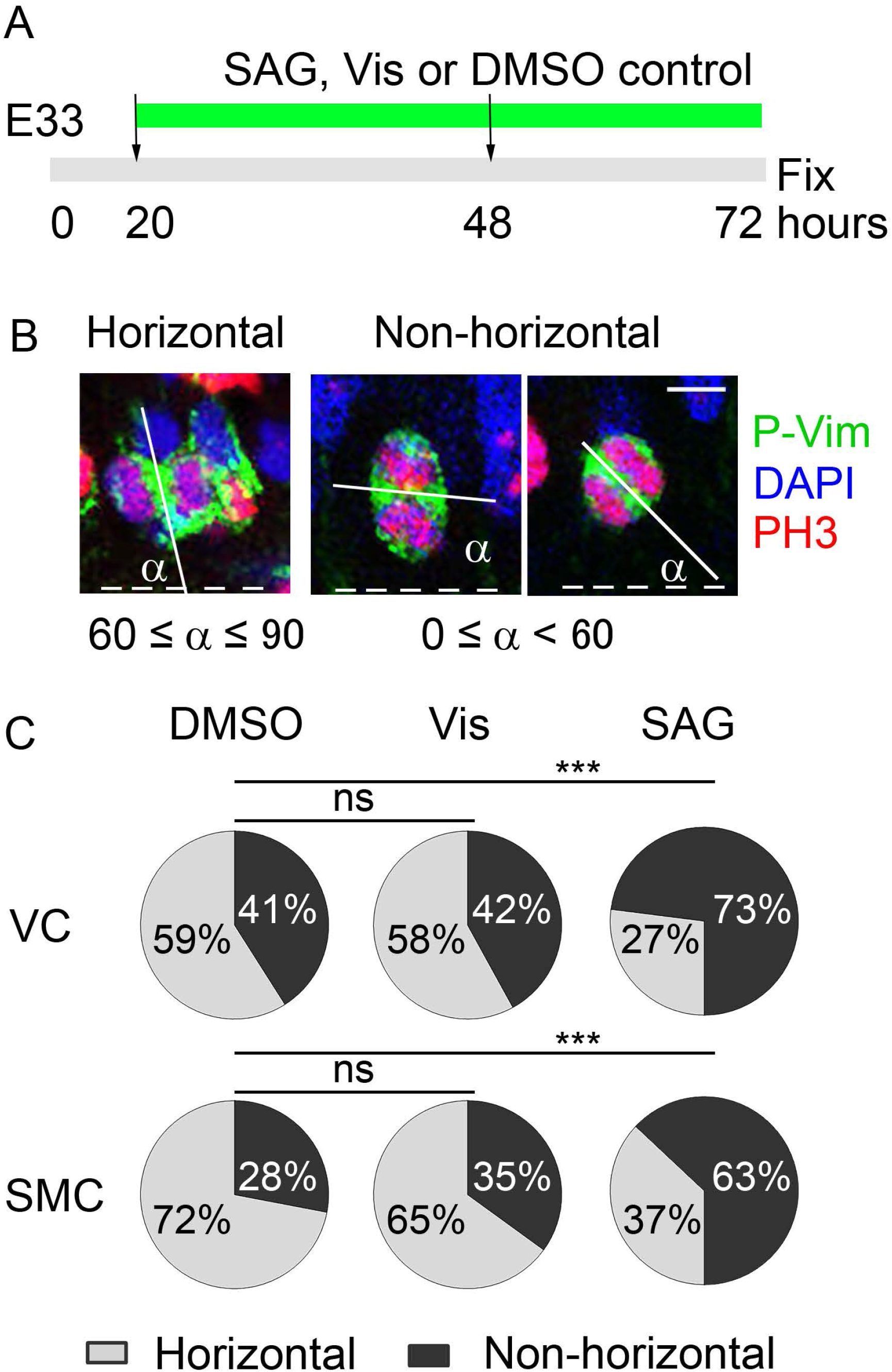
HH signaling is sufficient to promote non-horizontal division of vRGs *ex vivo*. (*A*) Diagram of the experimental scheme. Brain slices from E33 embryos were placed in culture and treated with SAG, vismodegib (Vis), or DMSO control for 2 days before fixation. (*B*) Micrographs of vRGs labeled with antibodies to P-Vim (green) and PH3 (red) and stained with DAPI (blue). Scale bar = 5 μm. The solid lines indicate the cleavage planes of anaphase cells, and the dotted lines indicate the ventricular surface. α indicates the acute angle between the solid and dotted lines. The modes of division are categorized as “horizontal” (60° < α ≤ 90°) or “non-horizontal” (0° ≤ α < 60°). (*C*) Quantification of acute angles (α) in the visual cortex (VC) and the sensorimotor cortex (SMC) in the slices placed in culture. Chi-square test: ns, *P* > 0.05; *** *P* < 0.001.

These *in vivo* and *ex vivo* results suggest that HH signaling promotes non-horizontal division of vRGs in a restricted cortical developmental stage in ferrets. Is this restriction unique to ferrets? To answer this question, we examined vRG division angles in mice. Previously, we showed that elevated HH signaling in *GFAP::Cre; SmoM2* mutant mice, in which SMOM2 (an active mutant form of SMO) constitutively activates HH signaling in vRGs, dramatically increased non-horizontal vRG divisions at E14 (to 42% in mutant mice vs. 15% in control mice, *P* < 0.01) (Wang et al. 2016). Remarkably, 1 day later at E15, non-horizontal vRG divisions decreased significantly to 24% in the mutant mice (Fig. 3D). Therefore, the effect of HH signaling on the vRG division angle was also dependent on the cortical developmental stage in mice.

### HH Signaling Maintains oRGs in the OSVZ to Expand Upper-Layer Neuron

After being produced from vRGs during a limited developmental window, oRGs self-renew in the OSVZ to maintain themselves (Martínez-Martínez et al. 2016). In the sensorimotor cortex at E35, even though the incidence of non-horizontal, oRG-producing divisions was minimal (Fig. 3), HH signaling was necessary and sufficient to expand oRGs (Figs. 1 and 2). In the visual cortex, SAG increased oRG numbers (Fig. 2) without affecting the vRG division angle (Fig. 3). These results might reflect the role of HH signaling in the phase of oRG expansion after their initial production from vRGs. To test whether HH signaling was required to maintain oRGs in the OSVZ in ferrets, we treated pregnant ferrets with vismodegib daily from E36 to E39, past the window for oRG production from vRGs (Martínez-Martínez et al. 2016), and collected the brains from ferret kits at postnatal day 2 (P2) (Fig. 5A). Vismodegib significantly reduced the number of oRGs (Figs. 5B and C and Supplementary Fig. 3), indicating that HH signaling is required to maintain oRGs in the OSVZ. Vismodegib also reduced the number of IPCs but not that of vRGs. These decreases in the numbers of oRGs and IPCs resulted in a decrease in Satb2-expressing neurons that reside superficial to Ctip2+ layer V neurons (Fig. 5D and E), indicating the critical roles of oRGs and IPCs in the expansion of upper-layer neurons.

**Fig. 5.**
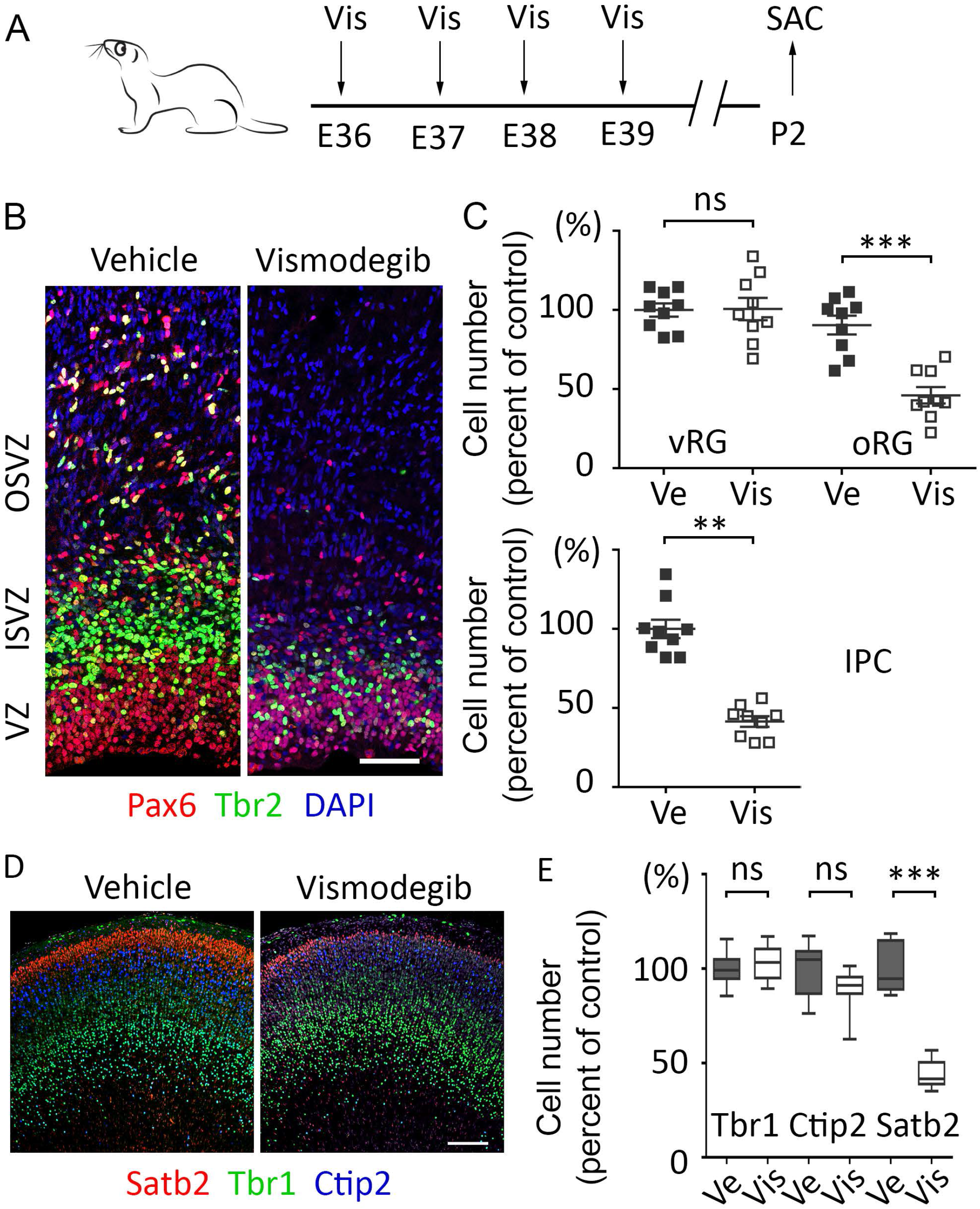
HH signaling is required for the expansion of oRGs, IPCs, and superficial-layer neurons. (*A*) Diagram of the treatment scheme. Pregnant ferrets were treated daily with vismodegib (Vis) (25 mg/kg) or vehicle (Ve) by oral gavage from E36 to E39. Brains were collected at postnatal day 2 (P2). (*B*) Micrograph of a P2 sensorimotor cortex labeled with antibodies to Pax6 (red) and Tbr2 (green) and stained with DAPI (blue). Scale bar = 50 µm. (*C*) Relative column densities of vRGs, oRGs, and IPCs. Mean ± SEM. (*D*) Micrograph of P2 sensorimotor cortex labeled with antibodies to Satb2 (red), Ctip2 (blue), and Tbr1 (green). (*E*) Box plots with min/max whiskers that show the relative densities of Satb2^+^ (layer II–IV), Ctip2^+^ (layer IV), and Tbr1^+^ (layer VI) neurons in the sensorimotor cortex. Mann–Whitney test: ns, *P* > 0.05; ** *P* < 0.01; *** *P* < 0.001.

To test directly whether HH signaling maintained oRGs by promoting their self-renewal, we prepared cultures of brain slices from E39 embryos, treated them with BrdU for 24 h, then chased the slices without BrdU for 32 h to identify the cells produced during the 24 h of BrdU treatment (Fig. 6A). In vehicle-treated controls, 38% of BrdU-labeled cells remained as oRGs in the OSVZ after 32 h of chase (Fig. 6B and C). SAG did not change the proportion of BrdU^+^ IPCs; however, it significantly decreased the proportion of BrdU^+^ Pax6^−^ Tbr2^−^ cells and increased the proportion of BrdU^+^ oRGs in the OSVZ to 62%, indicating that HH signaling promotes the self-renewal and maintenance of oRGs in the OSVZ in ferrets.

**Fig. 6.**
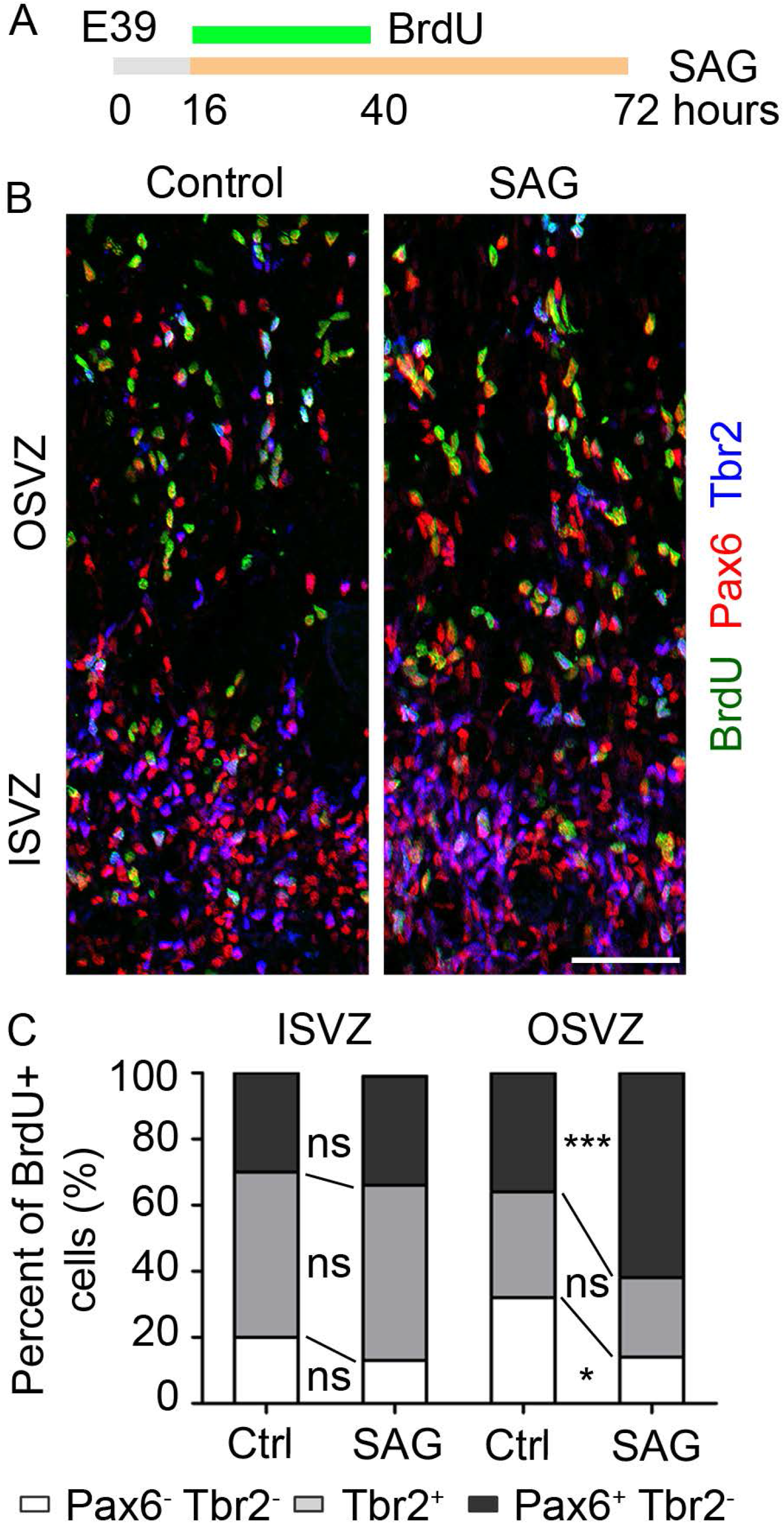
HH signaling promotes self-renewal of oRGs *ex vivo*. (*A*) Diagram of the experimental scheme. Brain slices from E39 embryos were placed in culture and treated with SAG and BrdU for the indicated periods before fixation. (*B*) Micrographs of the SVZ labeled with antibodies to Pax6 (red), Tbr2 (blue), and BrdU (green). Scale bar = 50 μm. (*C*) Quantification of the oRGs (BrdU^+^Pax6^+^Tbr2^-^ cells) and IPCs (BrdU^+^Tbr2^+^) as a percentage of the total BrdU^+^ cells in the OSVZ and the ISVZ. Mann–Whitney test: ns, *P* > 0.05; * *P* < 0.05; *** *P* < 0.001.

## Discussion

The relative expansion of the OSVZ, IPs, and oRGs in gyrencephalic species, as compared to those in lissencephalic species, have implicated them as playing crucial roles in neocortical development and evolution, triggering intense interest in the mechanisms underlying their expansion (Lui et al. 2011; Borrell and Götz 2014; Florio and Huttner 2014; Sun and Hevner 2014; Dehay et al. 2015; Vaid and Huttner 2020). Previously, we identified HH signaling as being necessary and sufficient to expand IPCs, oRGs, and upper-layer neurons in the mouse, a lissencephalic species (Wang et al. 2016). This was the first signaling pathway to be so identified. Here, we have shown that these functions of HH signaling are conserved in a gyrencephalic species, the ferret. Furthermore, we have shown that the cellular mechanisms by which HH signaling expands oRGs and IPCs are also conserved. HH signaling promotes proliferation of IPCs during embryonic development. It also promotes non-horizontal division of vRGs to produce oRGs in an early restricted phase before the peak of superficial-layer neuron production. Beyond this restricted phase, HH signaling promotes self-renewal of oRGs (Fig. 7).

**Fig. 7.**
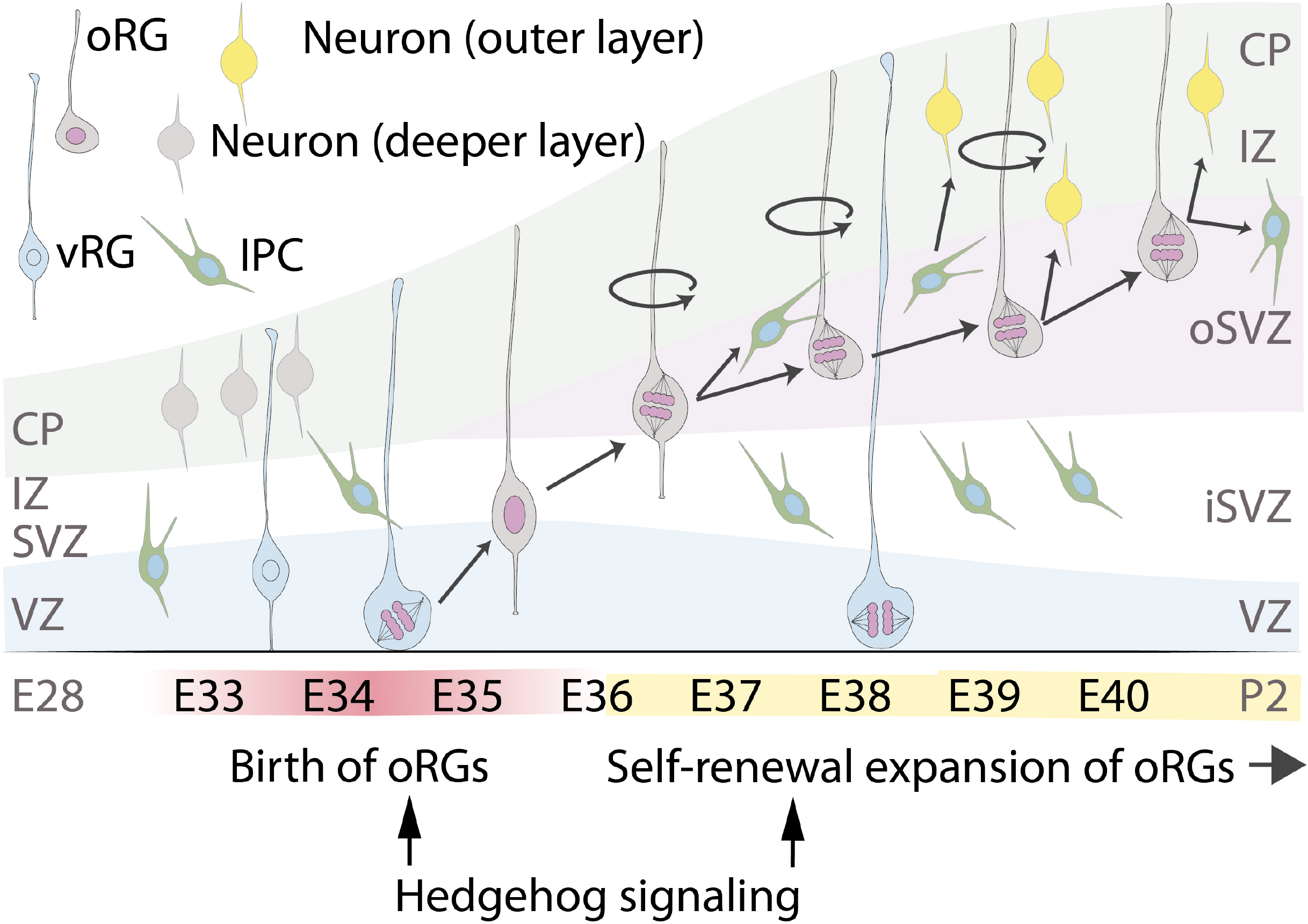
Biphasic function of HH signaling in the production and maintenance of oRGs: HH signaling promotes oRG production by regulating the spindle orientation of vRGs during the early restricted window and subsequently maintains the oRG population by promoting their self-renewal. A timeline of the development of the visual cortex is shown.

Ferrets are evolutionarily further from mice than are humans; therefore, the functions of HH signaling in NPCs are probably conserved in most mammalian lineages, including humans. Consistently, mutations in HH signaling components affect human brain size (Nanni et al. 1999; Heussler et al. 2002; Ginocchio et al. 2008; Derwińska et al. 2009; Twigg et al. 2016; Shiohama et al. 2017; Klein et al. 2019), and HH signaling is required for the expansion of oRGs in cerebral organoids (Wang et al. 2016). How, then, does the conserved function of HH signaling shape species-specific convoluted or smooth neocortices? The answer appears to lie, in part, in the different levels of HH signaling activities in different species. The expression levels of *Gli1*, a well-established readout of HH signaling activity, in the developing neocortex are higher in humans and ferrets than in mice, indicating that HH signaling activity is stronger in gyrencephalic species than in lissencephalic species (Wang et al. 2016; Matsumoto et al. 2020). Notably, Sonic hedgehog (SHH) protein levels in the developing neocortex are higher in ferrets than in mice, suggesting a potential mechanism for differential HH activities in ferrets and mice (Matsumoto et al. 2020). Major drivers of evolution are species-specific changes in regulatory elements of conserved essential genes. Regulatory elements of the *Shh* gene may vary between gyrencephalic and lissencephalic species, resulting in different SHH levels. Notably, a recent study showed that key components of the HH signaling pathway, including *SMO, GLI2*, and *GLI3*, are associated with human-specific regulatory elements, which are enriched in genes expressed in oRGs (Won et al. 2019). Together, these findings indicate that species-specific regulation of gene expression in oRGs might play key roles in neocortical expansion and that components of the HH signaling pathway are under such regulation, enabling the conserved HH signaling to shape the neocortex in a species-specific manner.

In addition to contributing to the evolution of divergent neocortices, differential HH signaling activities within a species might contribute to the differential growth of the neocortex, forming folds (gyri) and fissures (sulci). In ferrets, *Gli1* expression is higher in prospective gyri than in prospective sulci, indicating that HH signaling activities are higher in prospective gyri than in prospective sulci (de Juan Romero et al. 2015; Matsumoto et al. 2020). Therefore, our findings predict that the higher HH signaling activity in prospective gyri results in a local increase in the number of neurons and their dispersion, promoting gyrus formation.

In the lissencephalic mouse cortex, the frequency of non-horizontal division of vRGs is rather low (15%), which is consistent with the sparsity of oRGs in mice (Wang et al. 2016). The HH signaling gain-of-function mouse model (*GFAP::Cre; SmoM2*), which develops a convoluted neocortex, exhibits approximately three times more non-horizontal divisions when compared to wildtype mice (Wang et al. 2016). vRGs in human cerebral organoids display a high intrinsic level of non-horizontal divisions (54%), which is dependent on HH signaling (Wang et al. 2016). We have shown that the ferret visual cortex at the peak of oRG production resembles human organoids, with 62% of the vRGs dividing non-horizontally in an HH signaling–dependent manner. Therefore, a high proportion of non-horizontal vRG division appears to be characteristic of gyrated brains and ensures a large pool of oRGs for seeding to the OSVZ. Our data show that HH signaling is a conserved mechanism essential for this large seeding of oRGs.

Our findings further suggest that the effect of HH signaling on vRG division angle is spatiotemporally restricted during cortical development. In ferrets, HH signaling was required for the high level of non-horizontal vRG divisions in the visual cortex but not for the low level in the sensorimotor cortex at E35. This restriction is consistent with the findings of previous studies that showed that the production of oRGs destined for the OSVZ in the visual cortex is limited, with a peak at E34–E36 (Martínez-Martínez et al. 2016), and that the neocortex develops along a rostrocaudal gradient (Jackson *et al*. 1989; Noctor et al. 1997; Reillo and Borrell 2012). vRGs in the mouse also responded to HH signaling with a similar developmental restriction. Interestingly, in both ferrets and mice, the window in which HH signaling can affect the vRG division angle precedes the period of peak production of superficial-layer neurons, whose expansion coincides with the expansion of oRGs and the neocortex (Smart et al. 2002; Lukaszewicz et al. 2005; Fietz et al. 2010; Hansen *et al*. 2010; Reillo et al. 2011; Martínez-Cerdeño et al. 2012). These results suggest that the competence of vRGs to produce oRGs in response to HH signaling changes as neocortical development progresses, which may result in the separation of the vRG and oRG lineages. In humans, vRGs even change physically during the transition from deep to superficial neuron production (Nowakowski et al. 2016); after gestation week 16.5, the basal process of a vRG loses contact with the pial surface and terminates in the OSVZ. It will be interesting to investigate whether this physical change in human vRGs occurs gradually, following the spatial gradient of neocortical development, and whether it coincides with the ability of vRGs to respond to HH signaling and produce oRGs.

In addition to large-scale production of oRGs to expand the neuronal population, gyrencephalic species employ extended neurogenic time, which requires prolonged self-renewal of oRGs after their production during a restricted period. Our data show that HH signaling promotes oRG self-renewal. A recent study by Matsumoto et al. also showed that HH signaling enhances oRG self-renewal in ferrets; this was accomplished by introducing an activator and an inhibitor of HH signaling into E33 embryonic ferret cortices through *in utero* electroporation (Matsumoto et al. 2020). Therefore, two independent studies using different techniques have shown that HH signaling promotes the self-renewal of oRGs.

In summary, HH signaling has critical and conserved functions in expanding oRGs, IPCs, and upper-layer neurons, which together promote the development of gyrencephalic brains. These functions appear to be regulated species specifically, spatiotemporally, and cell-type specifically. Future investigations of such specific regulatory mechanisms and of the molecular mechanisms by which HH signaling affects the behaviors of vRGs, oRGs, and IPCs will provide important insights into neocortical evolution and development.

## Acknowledgments

We thank the staff of the Animal Resource Center, the Cell and Tissue Imaging Center, and the Veterinary Pathology Core at St. Jude Children’s Research Hospital for technical assistance. We thank Dr. Burgess Freeman for helping us to determine vismodegib and SAG doses for in vivo treatment. We thank Dr. Arnold Kriegstein’s laboratory for the protocols for ferret organotypic slice culture and viral transduction. We thank Drs. Xinwei Cao and Kristen Olesen for reviewing the manuscript. We thank Keith A. Laycock, PhD, ELS, for scientific editing of the manuscript. This work was supported by a Whitehall Foundation Research Grant, by the American Lebanese Syrian Associated Charities (ALSAC), and by the National Institutes of Health (R01NS100939 to Y.-G. H.), maintained by St. Jude Children’s Research Hospital. The content is solely the responsibility of the authors and does not necessarily represent the official views of the National Institutes of Health.

## Conflict of Interest

None declared.

## Figure Captions

**Supplementary Fig. 1.**The effectiveness of vismodegib treatment. (*A*) The cerebellum (left) and the cerebrum (right) of E35 embryos stained with H&E after treatment with vismodegib (25 mg/kg) or vehicle for 2 days (E33 and E34). (*B*) Blood vessels in the brain immunostained for vGLUT1. Scale bar = 100 μm. (*C*) Vismodegib decreased GNP proliferation in the external granular layer (EGL) of the cerebellum, as indicated by phospho-histone H3 labeling. Scale bar = 25 μm.

**Supplementary Fig. 2.**Vismodegib did not induce apoptosis in the cerebral cortex. (*A*) E35 cortices labeled for cleaved caspase 3 after treatment with vismodegib (Vis) (25 mg/kg) or vehicle (Ve) for 2 days (E33 and E34). (*B*) Quantification of the density of cleaved caspase 3–positive cells. Mean ± SEM. Mann–Whitney test: ns, *P* > 0.05.

**Supplementary Fig. 3.**HH signaling is required for the expansion of oRGs and IPCs.

(*A*) Micrograph of the P2 visual cortex (VC) labeled with antibodies to Pax6 (red) and Tbr2 (green) and stained with DAPI (blue) after daily treatment with vismodegib (Vis) (25 mg/kg) or vehicle (Ve) by oral gavage from E36 to E39. Scale bar = 50 μm. (*B*) Relative column densities of vRGs, oRGs, and IPCs. Mean ± SEM. Mann–Whitney test: ns, *P* > 0.05; ** *P* < 0.005; *** *P* < 0.001.

## Notes

### Competing Interest Statement

The authors have declared no competing interest.

