## Supplementary figures for "Biphasic roles of hedgehog signaling in the production and self-renewal of outer radial glia in the ferret cerebral cortex"

Fig. S1

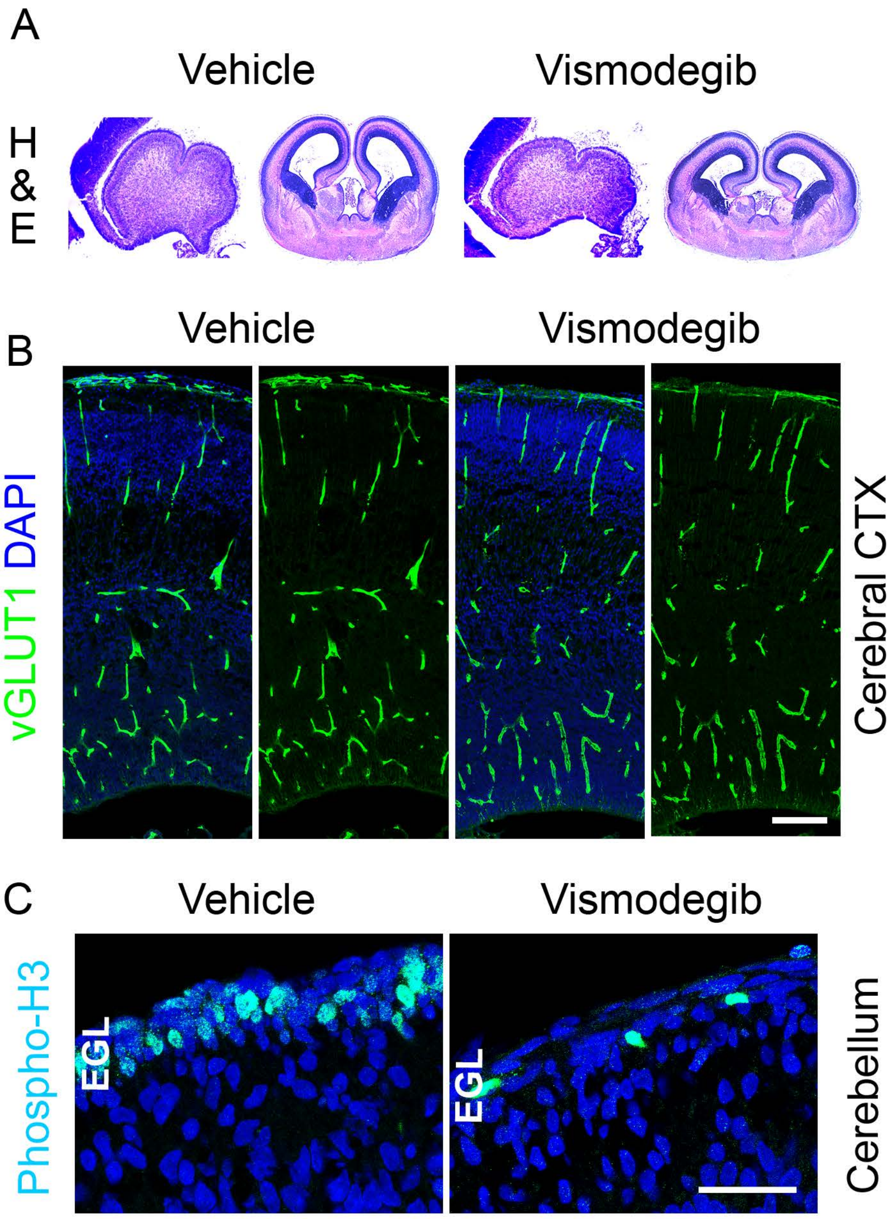


Fig. S1. The effectiveness of vismodegib treatment. (A) The cerebellum (left) and the cerebrum (right) of E35 embryos stained with H&E after treatment with vismodegib (25 mg/kg) or vehicle for 2 days (E33 and E34). (B) Blood vessels in the brain immunostained for vGLUT1. Scale bar = 100 μm. (C) Vismodegib decreased GNP proliferation in the external granular layer (EGL) of the cerebellum, as indicated by phospho-histone H3 labeling. Scale bar = 25 μm.

Fig. S2

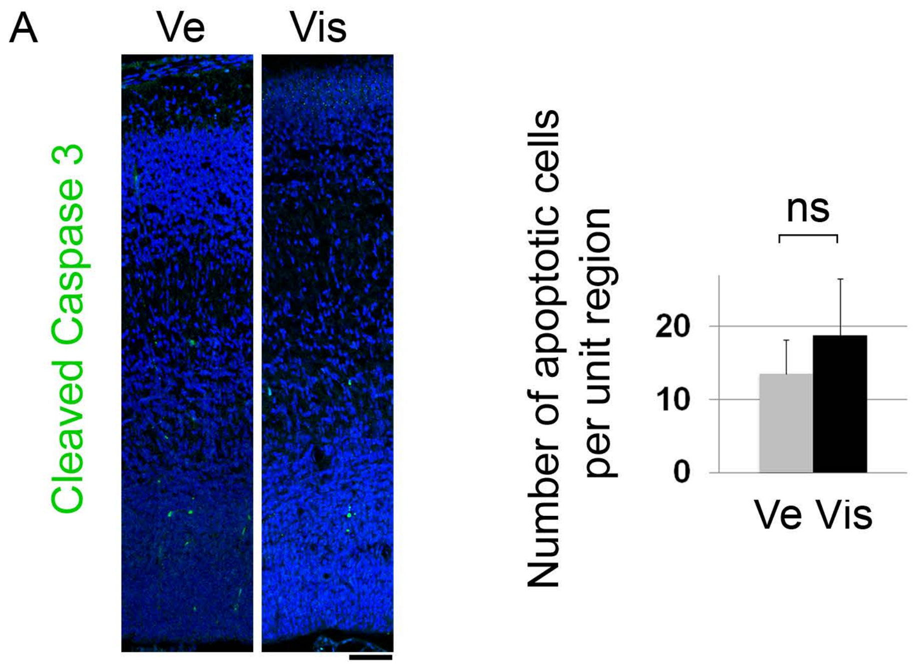


Fig. S2. Vismodegib did not induce apoptosis in the cerebral cortex. (A) E35 cortices labeled for cleaved caspase 3 after treatment with vismodegib (Vis) (25 mg/kg) or vehicle (Ve) for 2 days (E33 and E34). (B) Quantification of the density of cleaved caspase 3–positive cells. Mean ± SEM. Mann–Whitney test: ns, $P > 0.05$.

Fig. S3

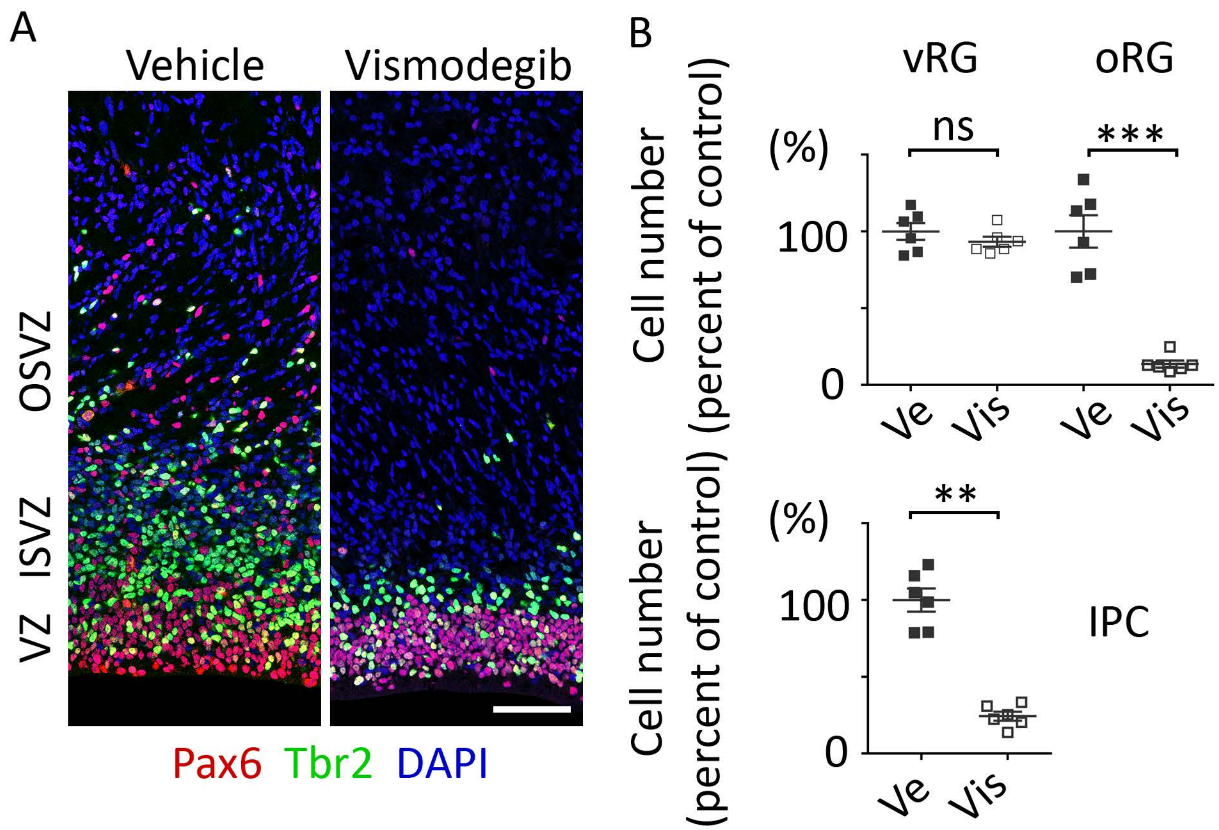


Fig. S3. HH signaling is required for the expansion of oRGs and IPCs. (A) Micrograph of the P2 visual cortex (VC) labeled with antibodies to Pax6 (red) and Tbr2 (green) and stained with DAPI (blue) after daily treatment with vismodegib (Vis) (25 mg/kg) or vehicle (Ve) by oral gavage from E36 to E39. Scale bar = 50 μm. (B) Relative column densities of vRGs, oRGs, and IPCs. Mean ± SEM. Mann–Whitney test: ns, $P > 0.05$; ** $P < 0.005$; *** $P < 0.001$.
